# Effect of relative social rank within a social hierarchy on neural activation in response to familiar or unfamiliar social signals

**DOI:** 10.1101/2020.08.25.267278

**Authors:** Won Lee, Hollie N. Dowd, Cyrus Nikain, Madeleine F. Dwortz, Eilene D. Yang, James P. Curley

**Affiliations:** Department of Psychology, Columbia University, New York, New York USA; School of Public Health, Yale University, New Haven, Connecticut USA; Department of Psychology, University of Texas, Austin, Texas USA

## Abstract

Competent social functioning of group-living species relies on the ability of individuals to detect and utilize conspecific social cues to guide behavior. Previous studies have identified numerous brain regions involved in processing these external cues, collectively referred to as the Social Decision-Making Network. However, how the brain encodes social information with respect to an individual’s social status has not been thoroughly examined. In mice, cues about an individual’s identity, including social status, are conveyed through urinary proteins. In this study, we assessed the neural cFos immunoreactivity in dominant and subordinate male mice exposed to familiar and unfamiliar dominant and subordinate male urine. The posteroventral medial amygdala was the only brain region that responded exclusively to dominant compared to subordinate male urine. In all other brain regions, including the VMH, PMv, and vlPAG, activity is modulated by a combination of odor familiarity and the social status of both the urine donor and the subject receiving the cue. We show that dominant subjects exhibit robust differential activity across different types of cues compared to subordinate subjects, suggesting that individuals perceive social cues differently depending on social experience. These data inform further investigation of neurobiological mechanisms underlying social-status related brain differences and behavior.

## Introduction

For social species living in groups, the ability to competently interact with others is vital for an individual’s survival and reproductive fitness^1^. Appropriate behavioral responses require accurate social decision making, whereby animals evaluate their external environment and integrate salient social information with their current internal physiological status as well as contextual and social memories^2–5^. Social decision making should be thought of as a reciprocal process between two or more individuals, with one individual’s behavioral output serving as input to another individual, that invokes rapid updating of information^3^. Brain circuits involved in social decision making that are conserved across social vertebrates constitute the Social Decision Making Network (SDMN)^5–8^. Studies in rodents have highlighted the role that subregions of the SDMN play in regulating social decisions in various social contexts including reproduction, parental care, aggression^3,9–13^. However, the functional role of the SDMN in the context of ongoing reciprocal group-level interactions has yet to be extensively explored.

A social dominance hierarchy is a common social context that necessitates constant social decision making. The establishment and maintenance of social hierarchies requires individuals to learn and remember their experience with each social partner in the group^14^ and dynamically express dominant or subordinate behavior towards animals of relatively lower or higher rank, respectively. Individuals must also remain vigilant to general social context changes, such as the presence or absence of high ranking individuals or alterations to group composition^15–19^. Despite these seemingly difficult cognitive requirements, numerous social species including fish, rats, mole rats, and mice are capable of rapidly organizing themselves into a hierarchy. Studies in primates, fish and rodents have recapitulated social hierarchies in the laboratory^17–19^. Such paradigms provide researchers with the unique opportunity to study the biological correlates of social behavior within an ethologically relevant context. Our lab has developed a paradigm that consistently generates autonomous hierarchies of 12 outbred CD-1 male mice^20,21^. We have shown that these mice exhibit flexible and context-appropriate social behaviors and adjust their behavioral response rapidly^15,22–24^. For example, individuals inhibit aggressive behavior in the presence of the alpha male^15^, but are able to quickly ascend in social rank when the alpha male is removed from the group^12,17,19^. Furthermore, we have shown that altering the general social context (e.g. by removing the alpha male), is associated with immediate early gene activation throughout SDMN nodes.

In mice, information about social context is often relayed through chemosensory cues present in urine. These cues can convey information relating to reproductive condition, dominance status, familiarity, and individual identity with exposure to these cues being shown to guide neural activity and behavior^25–27^. For instance, female mice show differential patterns of immediate early gene activation when exposed to dominant versus subordinate male urinary odors even when they have no previous experience with the urine donors^28^. Male mice scent mark over unfamiliar urinary odors more than familiar ones and show different patterns of immediate early gene expression when presented with familiar versus unfamiliar subjects^29,30^. There is also evidence across species that individuals respond differently to social cues depending upon individual factors such as their sex, hormonal state, past social experiences and social status^3,31^. For example, the rate of counter scent-marking over unfamiliar male urine is higher in dominant male mice than subordinates^32,33^. In cichlids, dominant male fish show a higher percentage of activated neurons when presented with food or female cues while subordinate males respond more sensitively towards dominant male odors^31^. These differential responses to social cues according to their own social status appear to be plastic adaptive responses. For instance, alpha males are required to distinguish between familiar and unfamiliar cues as they defend their territories from unknown conspecifics^34,35^. Conversely, mid-ranked or subordinate individuals should be able to recognize dominant or unfamiliar individuals and display attention or show fear-related responses to these individuals, while they should show less aversion towards social cues from familiar subordinate individuals. Indeed, there is strong evidence that subordinate animals pay more attention to the behavior and cues of dominant individuals than dominant individuals do to the behavior and cues of subordinates^15,36,37^.

In the present study, we investigate how an individual’s own internal state (social status) modulates the neural activity of their SDMN in response to olfactory cues of conspecifics of differing social status and familiarity. Specifically, we examine how differences in internal social state (dominant versus subordinate social status) are associated with activation of the immediate early gene (IEG) cFos in the brains of male mice in response to exposure to urinary odors. Subjects were exposed to one of four different cue types: familiar dominant, familiar subordinate, unfamiliar dominant or unfamiliar subordinate. Familiar urine donors originated from the same social hierarchies as the subject, whereas unfamiliar urine donors originated from different social hierarchies in separate housing systems. While we mainly focused on SDMN brain regions, we also examined cFos immunoreactivity in several other brain regions related to stress responsivity and those regions previously described to have high sensitivity to different social cues^12,28,30^.

## Results

### Social hierarchy characteristics

All 8 social groups of 12 male mice formed significantly linear social hierarchies (Supplementary **Fig. S1**). By the end of the group housing period, the median modified Landau’s h-value and interquartile range of 8 social groups (IQR) was 0.69 [0.63 - 0.71] (all p<0.01; see Supplementary **Table S1** for values for each cohort). Triangle transitivity was 0.88 [0.84 - 0.93] (all p<0.001) and directional consistency 0.89 [0.87 - 0.93] (all p<0.001). As noted in the methods (see **Fig. 1.** For the experimental design), urine samples were collected from group housing between days 7-10, after the establishment of social hierarchies. All social groups in this study established linear and stable social hierarchy by the 6th day of group housing (**Fig. 2**; median 4 days [2.8 - 5.3]).

**Figure 1:**
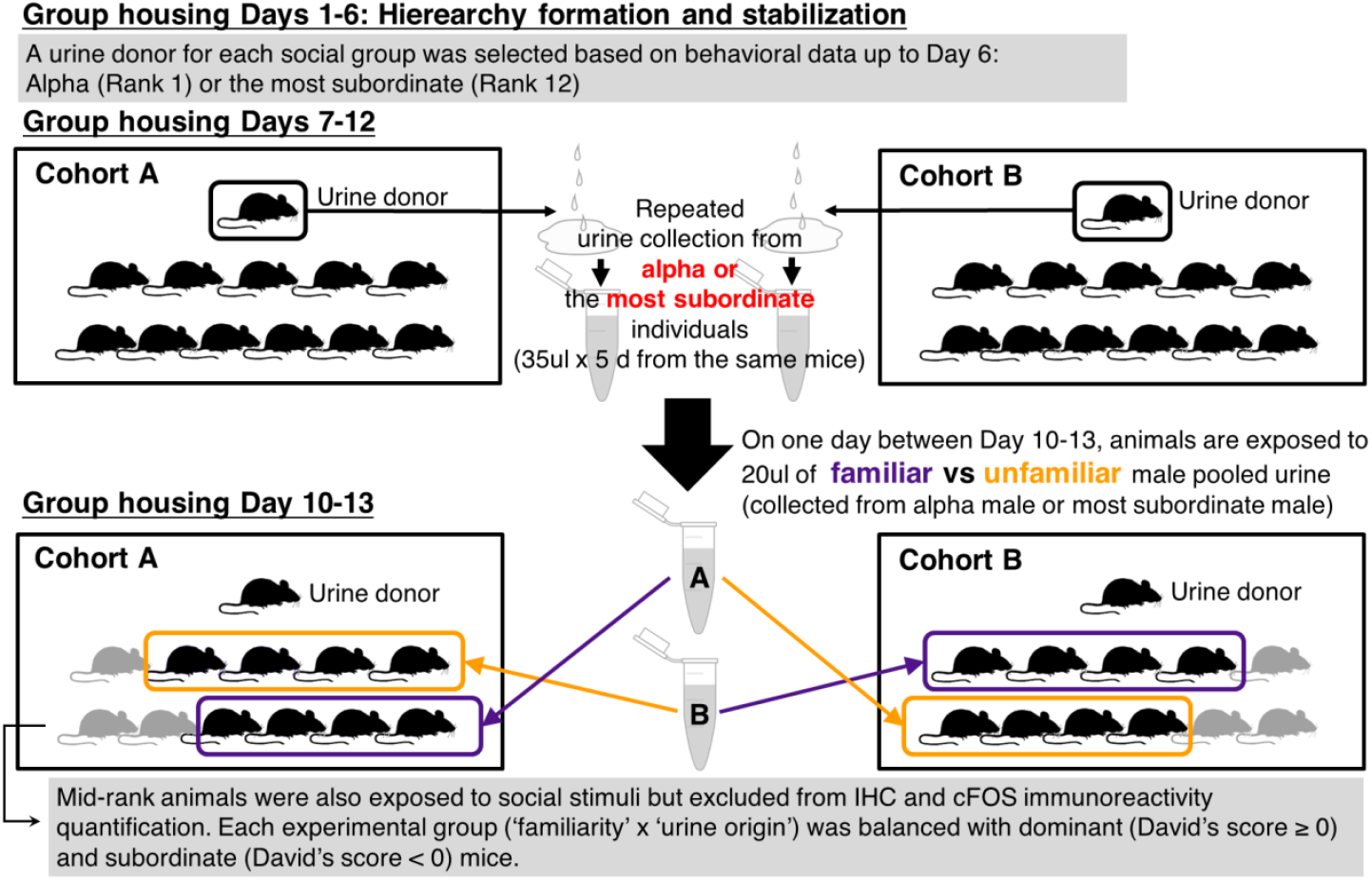
Schematic of the social stimulus exposure experimental design. 8 groups of 12 male CD-1 mice were housed in an ethologically relevant group housing system. All 8 groups formed linear social hierarchies by the start of urine collection from the donors (alpha (Rank 1) or the most subordinate (Rank 12) individual). On the last day of group housing (Day 10-13), all mice in social groups excluding the urine donor mouse were exposed to either familiar (purple) or unfamiliar (yellow) urine from either alpha or the most subordinate males.

**Figure 2:**
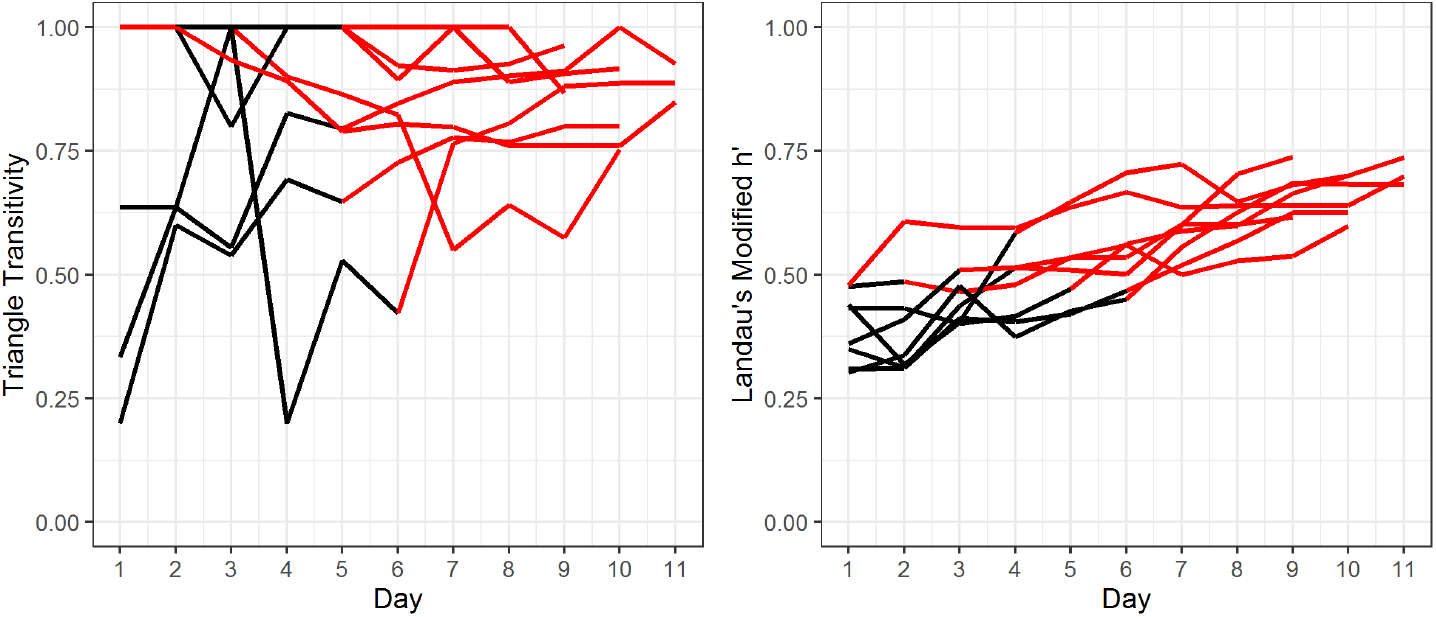
The triangle transitivity and Landau’s Modified h-value of each social group across the group housing period. All 8 cohorts formed linear and stable social hierarchies by day 6 of group housing. Each line represents a different social hierarchy. Red lines indicate significant values.

### Fos-immunoreactivity in response to social cues in dominant and subordinate male mice

For each brain region, we used Bayesian hierarchical modeling^38^ to estimate posteriors of 12 planned comparisons (**Table 1**, **Fig. 3**, Supplementary **Fig. S2**) to examine the effects of a) internal status - dominant vs. subordinate social status in the hierarchies; b) familiarity of social cues - familiar vs. unfamiliar social cues; c) the source of social cue - from the alpha or the most subordinate males in the hierarchies.

**Figure 3:**
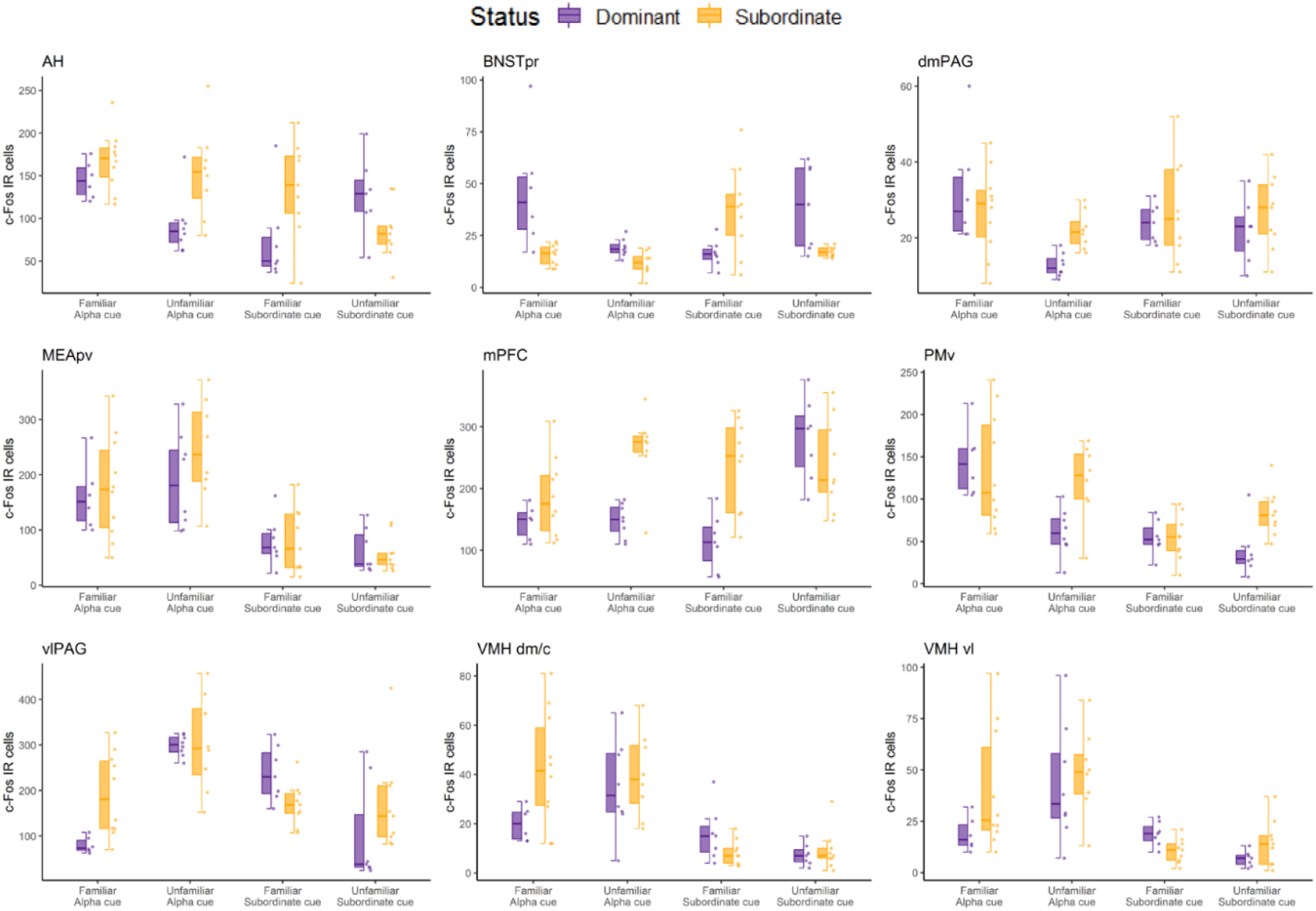
Fos-immunoreactivity in selected nine brain regions in response to urine from familiar alpha, unfamiliar alpha, familiar subordinate, unfamiliar subordinate males in dominant and subordinate male mice. Boxplots show median (horizontal bars), IQR (boxes) and 95% CI (whiskers). Raw data points of each group are also shown on the right side of each box plot. See Supplementary **Fig. S2** for the data of all 23 brain regions.

**Table 1:** Estimates and 95% highest density intervals (HDI) of 12 planned pairwise comparisons for cFos immunoreactivity in each brain regions. Bolded cells represent comparisons of which HDI did not overlap with 0.

| Region | Alpha urine |  | Sub urine |  | Alpha urine |  | Sub urine |  | Fam |  | Unfam |  |
| --- | --- | --- | --- | --- | --- | --- | --- | --- | --- | --- | --- | --- |
|  | Fam | Unfam | Fam | Unfam | Dom | Sub | Dom | Sub | Status: Dom | Sub | Dom | Sub |
|  | Internal state: Dominant status vs. subordinate status |  |  |  | Familiarity: Familiar cue vs. Unfamiliar cue |  |  |  | Source: Alpha urine cue vs. Sub urine cue |  |  |  |
| mPFC | -37 [-89, 21] | **-110 [-161, -53]\*** | **-114 [-174, -51]\*** | 34 [-30, 97] | -2 [-51, 50] | **-74 [-132, -17]\*** | **-142 [-211, -80]\*** | 4 [-59, 63] | 32 [-26, 86] | -50 [-109, 8] | **-117 [-175, -58]\*** | 22 [-34, 84] |
| NAcc | 0 [-14, 12] | -1 [-16, 10] | 0 [-27, 11] | 0 [-12, 25] | 0 [-10, 14] | 0 [-13, 14] | -1 [-28, 10] | 0 [-13, 2 4] | -1 [-22, 9] | -1 [-31, 6] | -14 [-35, 3] | -1 [-25, 8] |
| LSv | 0 [-69, 46] | -1 [-72, 38] | 0 [-55, 56] | 0 [-57, 56] | 0 [-51, 65] | 0 [-53, 53] | 0 [-51, 64] | 0 [-49, 61] | -2 [-105, 24] | -1 [-83, 28] | -2 [-99, 25] | -2 [-80, 32] |
| BNSTpr | 13 [-4, 35] | **7 [0, 12]\*** | -15 [-28, 2] | 15 [-3, 29] | 11 [-6, 32] | 3 [-3, 9] | -14 [-30, 3] | 14 [-2, 27] | 13 [-5, 35] | -17 [-29, 1] | -13 [-27, 4] | -5 [-9, 1] |
| mPOA | -14 [-33, 5] | 1 [-11, 29] | 0 [-19, 27] | 1 [-27, 59] | -6 [-30, 10] | 10 [-6, 32] | 0 [-48, 31] | 0 [-29, 30] | -12 [-42, 7] | 0 [-19, 19] | 0 [-59, 22] | -1 [-41, 16] |
| AH | -20 [-57, 15] | **-52 [-95, -12]\*** | **-57 [-103, -8]\*** | 36 [-3, 76] | **48 [8, 87]\*** | 16 [-23, 54] | **-48 [-93, -1]\*** | **47 [5, 85]\*** | **65 [17, 108]\*** | 27 [-12, 64] | -35 [-72, 8] | **57 [15, 97]\*** |
| MEApd | 0 [-14, 19] | 0 [-23, 14] | 0 [-32, 10] | 0 [-22, 20] | 0 [-17, 17] | 0 [-24, 12] | 0 [-28, 13] | 0 [-16, 24] | 0 [-12, 26] | 0 [-23, 12] | 0 [-19, 19] | 0 [-16, 24] |
| MEApv | -15 [-86, 59] | -50 [-128, 33] | -3 [-60, 58] | 6 [-45, 60] | -27 [-99, 52] | -60 [-134, 22] | 17 [-44, 73] | 25 [-28, 77] | **73 [5, 139]\*** | **86 [16, 150]\*** | **107 [39, 179]\*** | **165 [92, 235]\*** |
| BLA | 0 [-16, 18] | 0 [-25, 11] | 0 [-10, 28] | 0 [-10, 30] | 0 [-17, 19] | 0 [-24, 10] | 0 [-16, 22] | 0 [-12, 21] | 0 [-31, 10] | 0 [-17, 15] | 0 [-27, 11] | 0 [-9, 28] |
| CEA | 4 [-4, 14] | -4 [-11, 5] | 0 [-13, 16] | 6 [-10, 22] | 5 [-6, 14] | -4 [-11, 3] | 5 [-11, 24] | 10 [-3, 23] | -13 [-29, 2] | **-18 [-28, -5]\*** | -11 [-28, 2] | -2 [-13, 7] |
| VMH dm/c | **-18 [-33, -2]\*** | -4 [-21, 13] | 7 [-3, 17] | -2 [-9, 6] | -13 [-27, 2] | 1 [-16, 19] | 8 [-2, 18] | -1 [-8, 7] | 4 [-7, 15] | **29 [14, 44]\*** | **25 [11, 39]\*** | **28 [13, 41]\*** |
| VMHvl | -14 [-33, 4] | -8 [-29, 15] | **8 [0, 16]\*** | -8 [-17, 1] | -17 [-36, 3] | -8 [-31, 12] | **12 [5, 19]\*** | -4 [-13, 5] | -1 [-10, 10] | **22 [5, 40]\*** | **27 [9, 47]\*** | **29 [9, 46]\*** |
| CA1 | 0 [-25, 26] | -9 [-42, 11] | -1 [-31, 16] | 0 [-30, 24] | 0 [-25, 28] | -9 [-40, 10] | -12 [-42, 11] | -6 [-35, 16] | 17 [-7, 51] | 11 [-8, 37] | 1 [-20, 32] | 12 [-10, 45] |
| CA3 | 0 [-11, 8] | 0 [-7, 12] | 0 [-11, 8] | 0 [-11, 9] | 0 [-14, 6] | 0 [-9, 8] | 0 [-11, 10] | 0 [-10, 9] | 0 [-9, 11] | 0 [-7, 10] | 0 [-6, 15] | 0 [-8, 11] |
| DG | 0 [-8, 9] | 0 [-6, 12] | 0 [-13, 6] | 0 [-9, 10] | 0 [-12, 7] | 0 [-8, 8] | 0 [-14, 6] | 0 [-8, 9] | 0 [-6, 15] | 0 [-8, 9] | 0 [-6, 14] | 0 [-7, 10] |
| dIPAG | **-13 [-22, -4]\*** | **-17 [-31, -3]\*** | -2 [-15, 9] | -11 [-24, 1] | 2 [-7, 12] | -1 [-15, 11] | 7 [-4, 17] | -2 [-15, 12] | **-11 [-20, -1]\*** | 0 [-12, 11] | -6 [-17, 3] | -1 [-16, 14] |
| dmPAG | 1 [-8, 12] | **-9 [-14, -1]\*** | -2 [-10, 7] | -3 [-12, 4] | **13 [3, 23]\*** | 4 [-4, 11] | 2 [-6, 9] | 0 [-9, 9] | 3 [-6, 13] | 0 [-8, 10] | **-8 [-15, 0]\*** | -4 [-11, 4] |
| vlPAG | **-106 [-171, -43]\*** | 13 [-62, 95] | **64 [1, 126]\*** | -50 [-155, 52] | **-217 [-250, -174]\*** | **-90 [-188, -2]\*** | **128 [20, 217]\*** | 2 [-70, 77] | **-149 [-207, -89]\*** | 20 [-46, 88] | **185 [94, 272]\*** | **122 [17, 215]\*** |
| PMd | -10 [-37, 14] | 15 [-7, 43] | 5 [-11, 27] | 13 [-6, 32] | -14 [-47, 10] | 9 [-9, 34] | 10 [-11, 32] | 17 [-2, 33] | -11 [-40, 11] | 1 [-14, 25] | 11 [-9, 40] | 10 [-8, 28] |
| PMv | 12 [-34, 61] | **-56 [-91, -16]\*** | 3 [-30, 30] | **-40 [-75, -7]\*** | **75 [30, 112]\*** | 2 [-43, 48] | 16 [-19, 49] | -27 [-56, 3] | **78 [35, 117]\*** | **64 [21, 107]\*** | 22 [-13, 57] | 35 [-1, 71] |
| PVN | 0 [-11, 25] | 0 [-17, 19] | 0 [-20, 17] | 0 [-15, 23] | 0 [-22, 15] | 0 [-26, 11] | 0 [-18, 20] | 0 [-12, 26] | 0 [-23, 15] | 0 [-30, 8] | 0 [-18, 19] | 0 [-15, 22] |
| LH | **-47 [-82, -7]\*** | -27 [-57, 6] | -43 [-89, 11] | 17 [-23, 55] | -7 [-45, 27] | 11 [-19, 40] | -5 [-49, 31] | 47 [-4, 97] | -40 [-81, 4] | -31 [-78, 14] | -37 [-74, 0] | 1 [-31, 39] |
| DMH | 0 [-28, 13] | 0 [-27, 14] | 0 [-17, 25] | 0 [-13, 32] | 0 [-17, 23] | 0 [-17, 24] | 0 [-23, 20] | 0 [-16, 24] | 0 [-29, 15] | 0 [-16, 23] | 0 [-34, 10] | 0 [-17, 24] |

Among 23 analyzed regions, the MEApv was the only region for which all individuals responded with significantly higher cFos immunoactivity to alpha male urine compared to subordinate urine regardless of the subject’s own social status or the familiarity of the cue. In the VMHdm/c and VMHvl, alpha male urine induced significantly higher cFos immunoreactivity than subordinate urine for subordinate subjects exposed to familiar or unfamiliar urine, but only for dominant subjects when exposed to unfamiliar urine. Additionally, both dominant and subordinate animals showed a significant increase in cFos immunoreactivity in the PMv when exposed to familiar alpha urine compared to familiar subordinate urine but did not show any differences when exposed to unfamiliar alpha versus subordinate urine. Other brain regions showed context-dependent differences in cFos immunoreactivity when comparing exposure to alpha versus subordinate urine. For instance, both dominant and subordinate mice showed higher cFos immunoreactivity in the vlPAG when exposed to unfamiliar alpha urine compared to unfamiliar subordinate urine. Dominant mice also showed a decrease in cFos immunoreactivity in the vlPAG and dlPAG when exposed to familiar alpha urine compared to familiar subordinate urine. Two brain regions, the mPFC and dmPAG showed increases in cFos immunoreactivity only in dominant animals that were exposed to unfamiliar subordinate urine compared to unfamiliar alpha urine. In the AH, dominant mice increased cFos immunoreactivity to familiar alpha versus familiar subordinate urine, whereas subordinates showed increased cFos immunoreactivity to unfamiliar alpha versus unfamiliar subordinate urine.

When considering the familiarity of the urine cue, the vlPAG showed the biggest effect. Both dominant and subordinate mice showed significantly increased activity to unfamiliar alpha urine compared to familiar alpha urine. Dominant mice showed increased vlPAG cFos immunoreactivity in response to familiar subordinate compared to unfamiliar subordinate urine. Further, dominant, but not subordinate, mice exhibited increase in cFos immunoreactivity when exposed to familiar alpha compared to unfamiliar alpha urine in the PMv, dmPAG and AH. Dominant mice also showed increased activity to unfamiliar subordinate urine compared to familiar subordinate urine in the mPFC and AH, whereas they showed the opposite pattern in the VMHvl, responding significantly more to familiar subordinate urine than to unfamiliar subordinate urine. It is also worth noting that subordinate males responded with significantly increased cFos immunoreactivity to unfamiliar alpha urine compared to familiar alpha urine in the mPFC.

When comparing differences in the responses between dominant and subordinate animals to urine cues, effects were highly dependent upon the context of the social cue as already outlined above. Further, the strongest effects observed were that subordinate animals exhibited a more robust response to familiar alpha urine than dominant animals in the LH, vlPAG, and dlPAG. Subordinates responded more robustly than dominants to unfamiliar alpha urine in the PMv, dmPAG, dlPAG, AH and mPFC. When exposed to familiar subordinate urine, subordinates responded more than dominants in the mPFC and AH, but dominants responded more than subordinates in the VMHvl and vlPAG. Only one region, the PMv showed differential activity between dominant and subordinate mice exposed to unfamiliar subordinate urine, with subordinates showing significantly greater activity compared to dominants.

We next applied Flexible discriminant analysis (FDA) using data from all brain regions from all experimental conditions to classify similar patterns of brain activity into eight groups, and to identify which brain regions drove the strongest differences between groups. FDA classified the eight experimental groups correctly with seven discriminant functions. The first two functions explained 43.67% and 21.72% of the classification accuracy respectively (the corresponding spread of each data point is plotted in **Fig. 4**). Function 1 was mostly driven by mPFC (factor loading coefficient for Function 1: −1.53), MEApv (0.97), and AH (0.95). Function 1 appears to be primarily driven by being exposed to alpha urine cues compared to subordinate urine cues of any familiarity. Function 2 was loaded heavily with vlPAG (−1.52), AH (1.04), BNSTpr (0.73), VMHdm/c (−0.72). Function 2 was heavily driven by the difference between being exposed to familiar alpha or familiar subordinate urine in dominant individuals. The coefficient values for each brain regions for all seven functions are listed in Supplementary **Table S2**. The same analysis only including SBN nodes (AH, BNSTpr, dlPAG, dmPAG, LSv, mPOA, MEApd, MEApd, vlPAG, VMHvl, VMHdm/c) did not yield as accurate a prediction as when we included data from all brain regions (Supplementary **Fig. S3**).

**Figure 4:**
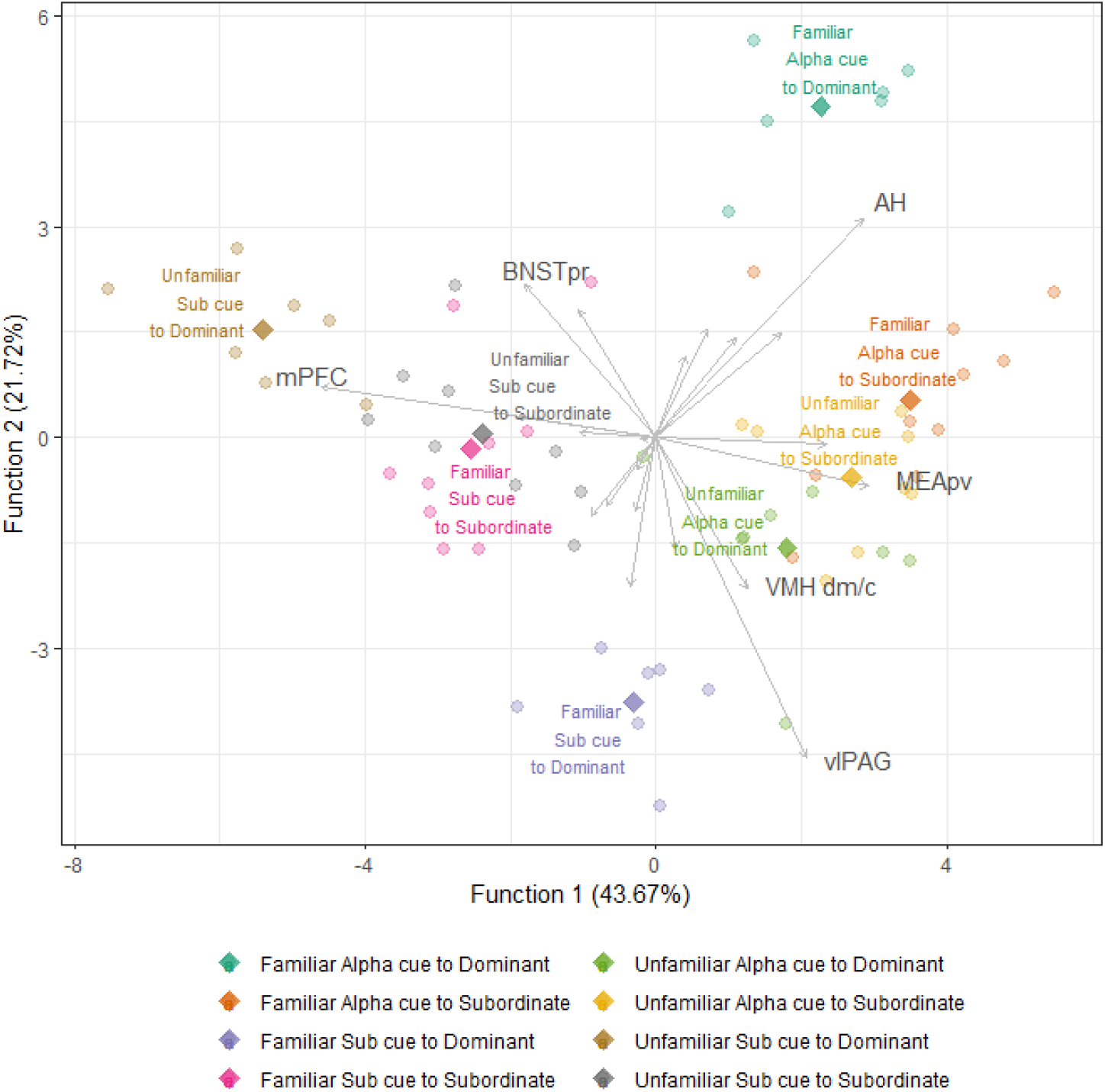
Flexible discriminant analysis of the eight experimental groups (status × familiarity × cue origin) as a function of cFos cell counts in all brain regions. Rhombi represent group centroids of each experimental group. Grey lines and represent the degree of separation (factor loading coefficient) each brain region contribute to Function 1 and Function 2. The longer the line is the larger contribution that brain has to for each function. For legibility of the entire plot, the grey lines were plotted with three times multiplication of the original (x, y) coordinates and only those brain regions that are most contributing to each function are labeled.

## Discussion

In this study we demonstrate that numerous brain regions exhibit cFos immunoreactivity in response to contextual features of social odor cues, depending on the internal state, or social status, of the subject processing the odor. We replicated our lab’s behavioral paradigm that generates linear social hierarchies within an ethologically relevant housing system. Groups of 12 outbred CD-1 mice were co-housed and allowed to establish stable linear social hierarchies and do so within a median of 4 days. Then with cFos immunohistochemistry, we identify excitatory activity patterns across 23 brain regions following exposure to a urinary odor cue. Induction of cFos protein expression is rapidly induced within neuronal nuclei upon depolarization of the cell membrane. Therefore, immunohistochemical detection of cFos reveals the location of excitatory activation in response to a given stimulus or event. It should be noted that this technique may not capture all activated neurons, as some neurons do not express the cFos gene, nor does it capture neuronal inhibition. Nevertheless, the major advantage of this technique is that it spatially maps activation of neural pathways among many regions simultaneously.

In total we examined cFos immunoreactivity following exposure to eight different cue conditions. In each condition, we varied three key features of the cue presentation: 1) the social status of urine donors; 2) the familiarity of the urine source; 3) the subject’s own internal state (social status). This results in eight different treatment groups (four social cue types for dominant and subordinate subjects). For each of the 23 brain regions we examined, we used pair-wise comparisons for each unique combination of these key features. While it is possible that some differences in cFos immunoreactivity between dominant and subordinate subjects may be due to underlying differences in stress susceptibility, we repeatedly habituated animals to the procedure to minimize this likelihood.

FDA analysis categorized subjects based on collective cFos immunorectivity in all 23 brain regions. We were able to classify subjects into each experimental group with 100% accuracy, indicating that collective brain activity from these brain regions is sufficient to determine which group each subject originated from. In other words, we are able to predict the social status of an individual and the type of odor cue they were exposed to based on the pattern of cFos immunoreactivity in the brain. The first two functions described by the FDA provide the strongest classification. Function 1 primarily discriminates between subjects that were exposed to urine from an alpha versus subordinate mouse, and mostly ignored familiarity and social status. Function 2 most efficiently discriminates between dominant subjects that were exposed to urine from a familiar alpha versus subordinate mouse. It should be noted that Function 2 does not separate subordinate subjects as efficiently. **Fig. 4**, which plots scores for Function 1 against Function 2 demonstrates that there is no overlap of dominant subjects across the four social cue conditions, whereas there is overlap between subordinate subjects exposed to each of the four social cue conditions. This suggests that brain-wide neural activation patterns induced by various social odors are more subtle in subordinate individuals and that discrimination of social cues relies on collective changes across the multiple brain regions. We hypothesize that Function 2 may represent level of saliency or the level of social vigilance towards given cues.

Notably, the FDA analysis is most successful when using data from all 23 brain regions rather than just brain regions associated with the SDNM. This demonstrates the importance of brain regions beyond this network for the differential processing of social cues from dominant and subordinate animals. However, five out of the six brain regions (BNSTpr, AH, MEApv, VMHdm/c, and vlPAG) that contribute most to the first two discrimination functions are also five of the six nodes of the social behavior network (SBN)^6,7^. The SBN in conjunction with the mesolimbic dopamine system comprises the SDMN and is considered to be an evolutionarily conserved set of interconnected brain regions whose coordinated activity drives social decision making and social behavior^5,7,39,40^. Consistent with our findings, other studies have also demonstrated that differential patterns of whole brain activity can be used to classify subjects during social discrimination tasks. For example, immediate early gene studies in African cichlid fish have also used FDA to identify differential patterns of neural activity throughout the SDMN in response to specific social contexts (e.g. exposure to dominant male, non-receptive female, aggressive gravid female, juveniles, etc.)^41,42^. Similarly, differential patterns of neural activity in the SDMN, olfactory bulb and mPFC can accurately predict whether male mice were exposed to novel or familiar juvenile mice or an empty cage^30^.

### Processing of Alpha versus Subordinate Olfactory Cues

Many animals, especially mice, use urinary odors to convey information about their identity, including social dominance^25,26,29,41,43^. The urine of male mice contains various chemosensory cues such as ESP1 (a pheromone shown to enhance female sexual receptivity) and major urinary proteins (MUPs)^44,45^. Dominant males show higher levels of MUPs, especially MUP20 (darcin)^27,46^. Our lab has previously shown that alpha CD-1 males living in social hierarchies of 12 mice produce higher levels of total MUPs and MUP20 (Lee et al., 2017). MUPs and MUP20 are used by dominant males to mark their territory and to signal dominance to other males, as well as to attract females^23,27,34,45^. Thus, the ability to distinguish between urine from a dominant versus a subordinate mouse is of high adaptive value in the context of territory marking, finding a mate and evaluating the social environment. Function 1 of the FDA analysis distinguishes whether the urine cue was from an alpha or subordinate mouse based on brain-wide cFos immunoreactivity patterns. This is predominantly driven by activity in the MEApv, mPFC, and AH. That is, these regions appear to be tuned to the dominance information conveyed by the urine cue. In our pairwise analysis, the MEApv is the only region to show differential activation based on the social status of the urine source alone, regardless of the familiarity with the source or the subject’s own social status. Specifically, this region shows greater activation in response to urine from alpha mice. The MEA receives olfactory information directly from both MOB and AON pathways^47^. Considering this region’s sensitivity to purely the dominance of the cue source, it is reasonable to hypothesize that it represents an early stage in an odor processing stream and is responsible for parsing out the most critical information first. In line with this idea, we also observe more refined responses to features of the cue presentation in downstream targets of the MEA, including the VMHvl and PMV. Prior studies have shown increased neural activity in MEApv neurons projecting to VMHdm in mice exposed to predator cat odor^48,49^. Indeed, it has been generally suggested that MEApv is involved in predator defense and MEApd is activated in the context of conspecific defense^50–52^. However, there is also evidence that the MEApv is activated when male mice are exposed to conspecific male odors^53^ or in social contexts with conspecifics such as mating and aggressive same-sex encounters^11^. It has been shown that the MEApv of female mice responds more robustly to dominant as opposed to subordinate urine from male conspecifics^28^, which is consistent with our results. Although we observed robust cFOS immunoreactivity to alpha urine, regardless of its familiarity or the social status of the subject animal, further investigation into neuronal type might reveal different response characteristics within this nucleus associated with these different social contexts further supporting a role for the MEApd in decision making in competitive social contexts^11,54,55^.

While the mPFC and AH also strongly contribute to Function 1, they also appear to have surprisingly refined responses to additional features of social cues, as revealed by the pairwise comparisons (**Table 1**). Importantly, the internal state of the subject (i.e. their own social status), highly influenced activation patterns. For instance, the mPFC of dominant subjects, but not subordinate subjects, exhibits increased activation to unfamiliar subordinate urine compared to unfamiliar alpha urine. Interestingly, this differentiation of subordinate versus alpha urine is not observed when the urine source is familiar. Furthermore, dominant subjects respond significantly more to unfamiliar subordinate urine than familiar subordinate urine. Previous studies corroborate our observation that social status is positively associated with activity in the mPFC^56,57^. The mPFC is also implicated in the processing of reward-based motivation, decisionmaking, cognition and fear and stress responses, all of which may be differentially regulated in dominant versus subordinate mice. Indeed, it has been shown that dominant mice exhibit greater motivation in social novelty exploration tasks^58^. In conjunction with our findings, it is possible that the mPFC is engaged in social motivation-related processes when presented with a novel social cue from a relatively subordinate source.

### Processing of Familiar versus Unfamiliar Olfactory Cues

As with distinguishing alpha versus subordinate urine, no one brain region appears to be exclusively tuned to familiar versus unfamiliar cues. The AH, which contributes heavily to Functions 1 and 2, exhibits differential activation across all three features of the cue presentation, with some activation patterns overlapping with those of the mPFC. While the AH serves important non-social functions (e.g. thermoregulation), it has also been implicated in social aggression, with elevated cFos immunoreactivity observed in mice selected for aggression^59^ as well as during offensive aggressive interactions in golden hamsters^60^. Interestingly, we find that the AH of dominant subjects responds more to familiar alpha urine than unfamiliar alpha urine. It is possible that AH activation is indicative of a response to a group-member with which they share a history of agonistic interactions. We must stress, however, that the AH exhibits differential activation in many pairwise comparisons, and therefore it’s response to social cues is likely to be influenced by many contextual factors.

Like the AH, the PMv of dominant subjects also exhibits increased cFOS immunoreactivity to familiar alpha urine compared to unfamiliar alpha urine. Both dominant and subordinate subjects exhibited increased activation of the PMv in response to familiar alpha versus familiar subordinate urine. Stimulating PMv neurons in male mice has been shown to trigger attack behavior towards conspecifics, whereas silencing them interrupts attacks^61^. Activation of the PMv can also induce general reward-related behaviors, including conditioned-place preference, and thus it has been suggested that it may play a role in organizing goal-oriented aggression^61^. That different olfactory cues activate the PMv of dominant and subordinate animals in our study may reflect differences in such goal-oriented aggression.

The PMv is innervated by monosynaptic reciprocal excitatory connections from the VMHvl^61,62^. Interestingly, in dominant subjects the differential responses of VMH regions contrast those of the PMv, such that dominant subjects exhibit increased VMHvl activation to unfamiliar alpha urine compared to unfamiliar subordinate urine. Subordinate subjects exhibit increased activation to alpha urine in general, regardless of familiarity. VMHvl also contributes heavily to Function 2, which we hypothesize represents level of saliency or level of social vigilance towards given cues. Neurons in VMHvl, especially those expressing estrogen receptor 1, show significant neural activation in aggressive, mating and defensive contexts^11,63–66^. These social contexts are highly salient, either aversive or appetitive, thus the sensitivity of VMHvl to these social contexts suggests that social information is gathered and the saliency of the contexts is estimated in VMH. Furthermore, optogenetic activation of VMHvl induces male mice to increase aggression^11^, whereas the activation of VMHdm/c induces defensive behavior^66^. These results suggest that VMH is significantly involved in decision-making of social behavior.

It has also been shown that the level of activation observed in the VMHvl is associated with differences in internal state (e.g. hunger or thirst), and thus activation could represent arousal-related information that guides social decision making^63^. The results from the current study also suggest that social status, a component of internal state, influences responses to social cue. While cFos immunoreactivity in the VMHvl of subordinate subjects increased in response to both familiar and unfamiliar alpha urine, it only increased in response to unfamiliar alpha urine in dominant subjects. This data suggests that the familiar alpha cue is not as salient to dominant subjects as subordinate subjects. While we’ve only discussed connections between the VMH, MEA and PMv, the VMH shares reciprocal connections with many brain regions including those in the SDMN^66–68^. The VMH’s abundant connections further supports the idea that it integrates various types of external social information with internal state.

The VMH, AH and PMd are sources of inputs to the PAG^50^, another region that responds significantly to urinary cues. Function 2 is most strongly driven by differences in the vlPAG which, like the VMH, exhibits increased activation in response to unfamiliar alpha urine in both dominant and subordinate subjects. Moreover, the vlPAG shows increased activation in dominant subjects when exposed to familiar subordinate urine compared to familiar alpha urine. In general, the PAG regulates the acquisition and expression of defensive behaviors in response to threatening stimuli and plays a role in fear learning^69^. For instance, optogenetic stimulation of the vlPAG leads to increased defensive behavior^70^ and micturition^71^, as well as facilitating Pavlovian fear conditioning^69^. In the current study, it is possible that increased activation of the vlPAG in response to an unfamiliar alpha reflects threat evaluation. In subordinate males, threat evaluation may lead to a defensive fear response, whereas in dominant subjects, it may elicit social vigilance and countermarking behavior. This would also explain dominant subjects’ increased activation in response to urine from a familiar subordinate source, as subjects would typically display vigilant dominant behavior towards them in a hierarchy context.

### Conclusion

Virtually all brain regions that respond to social cues in this study are modulated by social status. Dominant and subordinate subjects only share similar activity patterns across all conditions in the MEApv, where cFos immunoreactivity strongly predicted whether the cue source was an alpha or subordinate. Considering the MEApv’s early position in the olfactory processing stream, we hypothesize that this region encodes cue information that is immediately important for survival. Subcortical targets of the MEA and cortical regions also respond significantly to urinary cues, including but not limited to the mPFC, AH, PMv, VMH, and vlPAG. All of these regions exhibit differential activation across all three features of the cue presentation, meaning they are receptive to the effects of the cue source’s social status, familiarity with the source, and the subject’s internal state.

We believe these differences in neural activation across the brain could reflect the cognitive processes that enable these animals to behave in a socially plastic and contextually appropriate manner. The ability to extract relative social status information from social cues and to use that information to guide behavior is crucial for the maintenance of social hierarchies^1^. In addition to neural mechanisms that drive rapid behavioral flexibility from one context to the next, other types of plasticity occur on longer timescales and with varying reversibility^72^. In the current study, we show that cFos immunoreactivity in dominant subjects is highly differentiated between types of social cues, whereas differences in cFos immunoreactivity in subordinate subjects is more subtle across cues. This suggests that there are inherent differences in the processing of olfactory information related to dominant status. This hypothesis is supported by existing evidence that synaptic changes emerge in dominant individuals as they form dominance relationships^36,73^. These status-associated changes in neural plasticity can have implications across cognitive domains, such as learning and memory^74^. Further investigation into how reversible these status-related changes are will yield valuable insight to how social experience shapes the brain and life-history trajectories.

In conclusion, social information conveyed in odor cues is processed by a distributed network of regions, most of which respond to multiple contextual features. Rather than identifying distinct nodes where these features are encoded, this study demonstrates that the multifaceted nature of social cues is reflected in the distributed and nuanced way the brain processes them.

## Methods

### Subjects and housing

A total of 96 seven-week old male outbred Crl:CD-1 mice were obtained from Charles River Laboratories (Wilmington, MA, USA) and habituated to the facility in pairs in standard cages (27 × 17 × 12 cm^3^) with pine shaving bedding. Throughout the study, mice were housed under a 12:12 light:dark cycle (light cycle starting at 2400 hours and dark cycle starting at 1200hr) with constant room temperature (22-23°C) and humidity (30-50%) and provided with standard chow and water ad libitum. All mice were marked with a unique ID with non-toxic animal markers (Stoelting Co., Wood Dale, IL, USA). During pair housing, we determined dyadic dominant-subordinate relationships in each pair by focal observation during the dark phase of the light cycle for 2 hours per day for 3-4 days prior to group housing. Animals that consistently initiated and maintained aggressive behaviors were considered dominant and those that yielded to attacks and showed persistent subordinate behavior were considered to be subordinate. After 11-15 days of pair housing, each batch of 12 pairs (four batches total) were assigned to two separate social groups balancing for pair-housing social status. Each social group consisted of 6 randomly chosen dominant mice and 6 randomly chosen subordinate mice as previously described in Lee et al.,^23^. Mice were placed into custom built vivaria as previously described in So et al.^20^. Briefly, the vivarium (150 × 80 cm^2^ and 220 cm high; Mid-Atlantic, Hagerstown, MD, USA) consists of an upper level with three floor shelves and a lower level with five nest-boxes connected by tubes and all surfaces were covered with pine shavings (Supplementary **Fig. S4.**). The total surface of the vivarium is approximately 62,295 cm^2^, providing approximately 5191.25 cm^2^ per mouse. Standard chow (LabDiet 5001, PMI, St. Louis, MO, USA) and water were provided ad libitum at the top left and right side of the vivarium. Each social group consisted of six dominant males and six subordinate males based on pair housing observations. Live behavioral observations started on the first day of group housing and continued for 10-13 days per social group with an average of 22 hours of total observation evenly distributed across the housing period. During these observations, trained observers recorded all interactions of fighting, chasing, mounting, subordinate posture and induce-flee behaviors that occurred between any two individuals as further described in Williamson et al.^21^. All experiments were approved by the Columbia University Institutional Animal Care and Use Committee (IACUC Protocol No. AC-AAAP5405) and were conducted in accordance with National Health Institute (NIH; Bethesda, MD, USA) and Columbia University IACUC animal care guidelines.

### Urine collection

Based on the behavioral observation data up to Group housing Day 6, we selected a urine donor mouse from each social group (either alpha or the most subordinate mouse in the group) (see **Fig. 1**). On group housing Days 7-12, we collected the minimum volume of 35ul urine daily and repeatedly from the same urine donor. Each day, mice were quickly removed from the group housing environment and scruffed over Eppendorf tubes as described in Supplemental Methods in Lee et al.^23^. In one of eight social groups we used, we had difficulty collecting urine from a subordinate urine donor (subordinate mice often inhibit urination)^23,75^. Therefore, we housed the social group one more day in the group housing system to collect sufficient urine. Collected urine samples were immediately stored at −20°C until the day of social stimulus exposure. Urine samples were only defrosted once on the day of social stimulus exposure and pooled for each individual to minimize possible daily variation in urine contents.

### Social stimulus exposure

To minimize cFos immunoreactivity due to potential stress responses due to handling, we habituated mice to scruffing and urine exposure for the last 5 days of pair housing by applying 20ul of nuclease free water directly on their oronasal groove, following the procedure previously described in Martel and Baum^76^. On the last day of group housing (group housing day 10-13), all mice were removed from the vivaria simultaneously at zeitgeber hour 12 (start of the dark cycle). Then mice were isolated for 60-140 minutes before the social stimulus exposure in a standard cage filled with pine shaving and provided with ad libitum standard chow and water. After the habituation, all mice except the urine donor in each social group were gently scruffed and 20ul of urine was applied onto the oronasal grove of each mouse. Each mouse was then immediately placed back to its isolation cage, and 90 minutes after the exposure, was perfused transcardially. Mice were randomly assigned to two experimental groups: a) familiar urine exposure (the urine samples were collected from either alpha or the most subordinate in the same group as subjects), b) unfamiliar urine exposure (the urine samples were collected from either alpha or most subordinate individuals in different social group). A total of 64 mice from 8 social groups were used to visualize and quantify cFOS immunoreactivity in response to urine stimuli (familiar alpha male urine: 6 dominant mice and 10 subordinate mice; unfamiliar alpha male urine: 8 dominant mice and 8 subordinate mice; familiar subordinate urine: 7 dominant mice and 9 subordinate mice; unfamiliar subordinate urine: 7 dominant mice and 9 subordinate mice).

### Fos protein immunohistochemistry

Ninety minutes after the exposure to urine stimulus, mice were deeply anesthetized with mixture of ketamine and xylazine and perfused transcardially with pH7.4 0.1M phosphate-buffered saline (PBS) followed by 4% paraformaldehyde (PFA) in 0.1M PBS. Brains were removed and immersed overnight in 4% PFA then 30% sucrose in 0.1M PBS for at least 24 hours (4°C). Whole brains were sectioned coronally at 40um thickness in a cryostat and stored in 0.1 M PB azide. Four series of sections were taken, and every other series was used for cFOS immunostaining (80um between stained sections). Immunostaining was conducted according to free-floating avidin-biotin procedure, using the Vectastain ABC Elite peroxidase rabbit IgG kit (Vector Laboratories, Burlingame, CA, USA). Sections were first washed in 0.1 M PB for five minutes three times then incubated with 3% H2O2 for 5 minutes followed by three times of 5minute washes in 0.1M PB/0.1% Triton-X (PBT). The sections were incubated with 1% normal goat serum (NGS) in PBT for an hour followed by 4°C overnight incubation in primary Fos rabbit polyclonal IgG (sc-52; Santa Cruz Biotechnology, Inc. Dallas, Texas, USA) at a concentration of 1:5000 in 1 % NGS/PBT. Slices were washed three times in PBT for 5 minutes then incubated in biotinylated anti-rabbit IgG (Vectastain, Vector Laboratories) at a concentration of 1:200 in PBS. The sections were rinsed again three times with PBT for 5 minutes then incubated in avidin-biotin-peroxidase complex at a concentration of 1:250 in PBS for an hour. Sections were washed again with 0.1 M PBS three time for 5 minutes. H2O2 (final concentration as 0.02 %) was added to 0.02 % 3,3’-diaminobenzidine (DAB) solution just prior to incubation. Sections were incubated in DAB solution to visualize immunoreactivity for 2-4 minutes and the reaction was immediately stopped by washing sections in PBS for 1 minute for the first wash and 5 minutes for the remaining three washes. All sections were mounted on FisherBrand Plus slides within 24 hours then processed for serial dehydration followed by coverslipping with Krystalon mounting medium (Millipore Billerica, MA, USA).

### Quantification of Fos-immunoreactive cells

Images were obtained with a Leica DMi8 microscope (Leica Microsystems, Wetzlar, Germany) under a 10x objective at a magnification x100. Regions of interest were located using the mouse brain atlas of Paxinos and Franklin^77^. A total of 23 brain regions were analyzed and their bregmas are described in Supplementary Table. S1: anterior hypothalamus (AH), basolateral amygdaloid nucleus (BLA), principal nucleus of the bed nucleus of the stria terminalis (BNSTpr), field CA1 of hippocampus (CA1), field CA3 of hippocampus (CA3), central amygdaloid nucleus (CEA), dentate gyrus (DG), dorsolateral periaqueductal gray (dlPAG), dorsomedial hypothalamus (DMH), dorsomedial periaqueductal gray (dmPAG), lateral hypothalamus (LH), ventral lateral septal nucleus (LSv), medial amygdaloid nucleus, posterodorsal part (MEApd), medial amygdaloid nucleus, posteroventral part (MEApv), infralimbic/prelimbic regions of the prefrontal cortex (mPFC), medial preoptic area (mPOA), nucleus accumbens core (NAcc), dorsal premammillary nucleus (PMd), ventral premammillary nucleus (PMv), paraventricular nucleus (PVN), ventrolateral periaqueductal gray (vlPAG), ventromedial hypothalamic nucleus, dorsomedial central part (VMHdm/c) and ventromedial hypothalamic nucleus, ventrolateral part (VMHvl) (see Supplementary **Fig. S5.** for example images). For each brain regions, 3 brain sections per mouse were imaged and cell counts were summed. We overlaid images from The Mouse Brain in Steretaxic Coordinates^77^ to include only the exact portion of the region. We used Fiji^78^ to count cFos positive cells.

### Statistical analysis

All statistical analyses were undertaken in R version 3.6.4.^79^.

### Group dominance structure, emergence of hierarchies and individual social status

For each cohort, we aggregated the total number of wins and losses by each mouse from all recorded aggressive interaction over the entire group housing period into win/loss frequency sociomatrices. To confirm the presence of a linear social hierarchy in each social cohort, we calculated Landau’s h-value, triangle transitivity (ttri) directional consistency (DC) and associated p-values from 10,000 Monte-Carlo randomizations^80,81^ using compete R package^82^ as previously described in Williamson et al.^21^. Further, we calculated ttri and Landau’s h-value and their significance for aggregated win/loss sociomatrices each day of group housing to determine how quickly each cohort established a linear social hierarchy. Despotism of each alpha was derived by calculating the proportion of wins by the alpha to all aggressive interactions observed during group housing period. Emergence of linear and stable social hierarchies were additionally confirmed by calculating individual Glicko ratings and visualizing them for each cohort using PlayerRatings R package^83^. David’s score, a win proportion measure adjusted for opponent’s relative dominance in the hierarchy^84^, was calculated for each individual to assign individual social rank in each hierarchy using the compete R package^82^. Based on our observation of social hierarchy dynamics, we first categorized male mice living in social hierarchies into three distinct social status groups: alpha (rank 1, the highest David’s score), subdominant (David’s score >=0), and subordinate (David’s score <0; see Supplementary **Fig. S6.** for the David’s score of each individual in the study) as previously described in Lee et al.^23,85,86^. Based on David’s scores (on group housing Day 6), we selected two urine donors per cohort (alpha = the highest David’s score in the group, the most subordinate = the lowest David’s score). Alpha males were included for cFOS quantification only when they were not urine donors. We tested and did not find any statistical differences between alpha males and other dominant males in their cFOS immunoreactivity, thus we consolidated cFOS immunoreactivity data from alpha and other dominant (DS>0) together and refer them as ‘dominant’ throughout the manuscript.

### cFos immunoreactivity count data analysis

We used Bayesian hierarchical generalized linear modeling^38^ fitted in STAN^87^ with the brms R package^88^. We took advantage of this Bayesian approach as it can handle outliers in the data by implementing heavy-tailed distributions while accommodating heterogeneity of variances across groups^38^. Similar to traditional frequentist analysis of variance (ANOVA), but free from frequentist ANOVA’s strict assumptions, we conducted 12 planned pairwise comparisons among experimental groups (main effects; familiarity of the given social cue and individual social rank) for each brain region. We used flexible discriminant analysis (FDA) to examine whether cFos immunoreactivity across 23 brain regions can predict which of the eight experimental groups (status × familiarity × cue source) each individual corresponded to. We used the mda R package to conduct FDA^89^. As an extension of linear discriminant analysis (LDA), FDA is capable of handling non-normality or non-linearity among variables, thus FDA results in more robust classification of groups given the data compared to LDA when data violates the assumptions of LDA^90^.

## Acknowledgements

We thank Dr. Frances Champagne for suggestions in writing the manuscript and Curley Lab members for help with behavioral observations.

## Author contributions

W.L. and J.P.C. conceived and planned the experiments. W.L., H.N.D, and C.N. carried out the behavioral work. W.L., H.N.D, C.N, and E.D.Y performed immunohistochemistry and collected cFOS immunoreactivity count data. W.L. and J.P.C. analyzed the data and made figures in the manuscript. W.L., M.D and J.P.C. wrote the paper. All authors contributed to the final version of the manuscript.

