## Supplemental Materials for "Effect of relative social rank within a social hierarchy on neural activation in response to familiar or unfamiliar social signals"

2 – Department of Chronic Disease Epidemiology, Yale School of Public Health, New Haven, Connecticut USA

**SUPPLEMENTARY MATERIALS**

**Supplementary Table 1. Social hierarchy measures of each social group (A – H).** p-values for each Landau's *h'*, triangle transitivity (*ttri*), and directional consistency (DC) were all below 0.01.

| Cohort ID | Landau's *h'* | *ttri* | DC | despotism |
|---|---|---|---|---|
| A | 0.74 | 0.96 | 0.92 | 0.34 |
| B | 0.6 | 0.75 | 0.88 | 0.47 |
| C | 0.7 | 0.92 | 0.94 | 0.56 |
| D | 0.61 | 0.87 | 0.88 | 0.64 |
| E | 0.63 | 0.8 | 0.94 | 0.64 |
| F | 0.68 | 0.89 | 0.84 | 0.57 |
| G | 0.79 | 0.94 | 0.9 | 0.68 |
| H | 0.7 | 0.85 | 0.84 | 0.61 |

**Supplementary Table 2. Summary of flexible discriminant function analysis (FDA).**

| Region | V1 | V2 | V3 | V4 | V5 | V6 | V7 |
|---|---|---|---|---|---|---|---|
| **Percent explained** | **43.7** | **65.4** | **77.1** | **86.6** | **95.6** | **98.3** | **100** |
| **AH** | **0.95** | **1.04** | 0.55 | 0.47 | -0.41 | -0.03 | 0.16 |
| **BLA** | -0.09 | -0.15 | 0.03 | 0.12 | 0.03 | 0.18 | -0.54 |
| **BNSTpr** | -0.6 | **0.73** | 0.74 | -0.57 | -0.76 | -0.53 | -0.39 |
| **CA1** | 0.23 | 0.51 | 0.12 | -0.35 | 0.17 | -0.12 | -0.1 |
| **CA3** | 0 | 0.03 | -0.04 | 0.02 | -0.16 | 0.29 | 0.66 |
| **CEA** | -0.09 | -0.35 | 0.02 | 0.06 | -0.47 | -0.37 | -0.34 |
| **DG** | 0.36 | 0.47 | -0.93 | -0.88 | 0.49 | 0.09 | 0.09 |
| **dlPAG** | -0.3 | -0.37 | -0.66 | 0.42 | 0.37 | -0.34 | -0.13 |
| **DMH** | -0.01 | 0 | -0.15 | -0.21 | -0.09 | 0.57 | -0.49 |
| **dmPAG** | 0.78 | -0.03 | -0.8 | 0.54 | 0.01 | 0.3 | 0.23 |
| **LH** | 0.09 | -0.53 | 1.46 | 1.51 | -0.4 | -0.18 | 0.2 |
| **LSv** | -0.07 | 0 | -0.55 | -0.47 | 0.39 | 0.18 | -0.07 |
| **MEApd** | -0.35 | 0.6 | 0.05 | 0.23 | 0.29 | -0.41 | 0.65 |
| **MEApv** | **0.97** | -0.23 | 1.04 | 0.63 | -0.57 | -0.48 | -0.7 |
| **mPFC** | **-1.53** | 0.24 | 0.68 | -0.48 | 1.36 | 0.01 | -0.08 |
| **mPOA** | -0.64 | 0.1 | 0.51 | -0.19 | 0.14 | 0.77 | -0.14 |
| **NAcc** | -0.12 | -0.7 | -0.34 | 0.52 | -0.19 | -0.06 | 0.21 |
| **PMd** | -0.23 | -0.33 | 0.22 | 0.11 | -0.18 | 0.2 | 0.6 |
| **PMv** | 0.57 | 0.5 | -1.09 | 0.23 | 0.29 | 0.12 | -0.26 |
| **PVN** | -0.35 | 0.03 | -0.85 | -0.84 | -0.29 | -0.29 | -0.01 |
| **vlPAG** | 0.69 | **-1.52** | -0.01 | -0.27 | -0.03 | -0.18 | -0.31 |
| **`VMH dm/c`** | 0.42 | **-0.72** | 0.76 | -0.28 | 0.1 | 0.84 | 0.03 |
| **`VMH vl`** | 0.14 | 0.38 | 0.03 | -0.33 | 0.35 | -0.52 | 0.3 |

**Supplementary Fig. S1. Row 1 and 2)** Raw and binarized sociomatrices of wins and losses for all aggressive behaviors across all cohort (A-H). Each value represents the total number of wins by the individual in each row against the individual in each column for each behavior. The degree of redness represents the frequency of wins. Individuals are ordered by David's score. **Row3)** Temporal dynamics of individual Glicko ratings throughout entire group housing period.

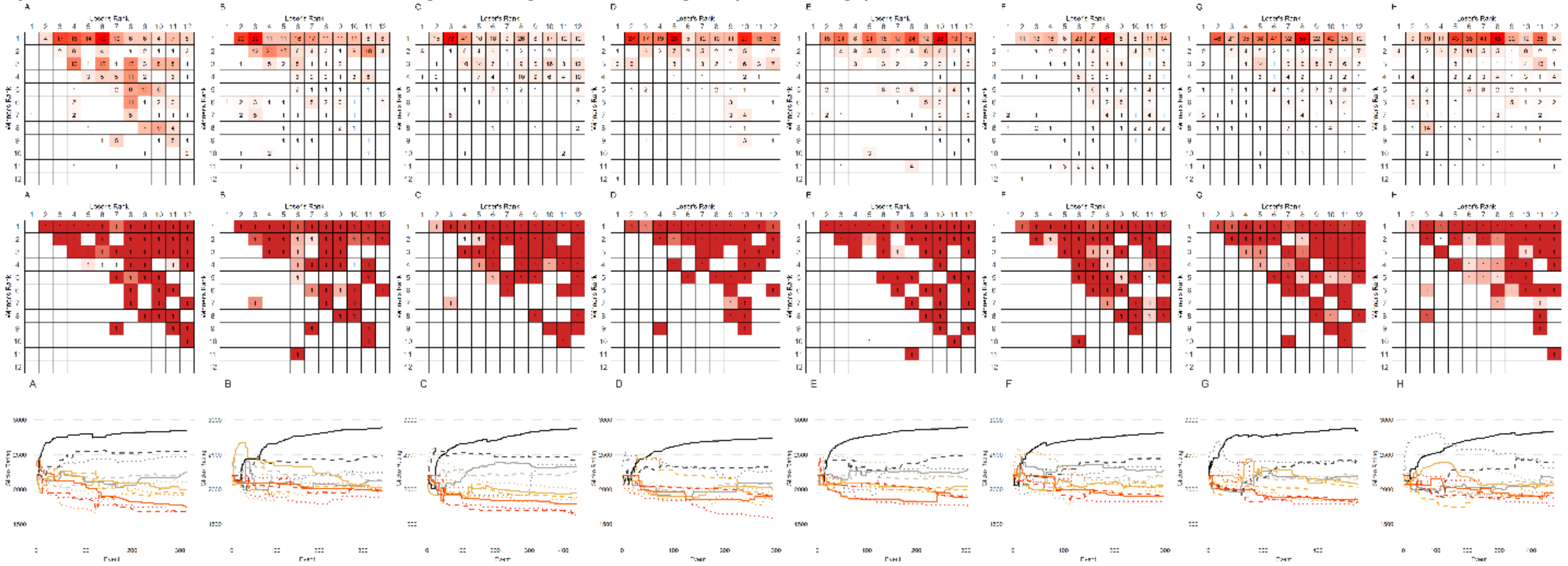

**Supplementary Fig. S2. Fos-immunoreactivity in response to urine from familiar alpha, unfamiliar alpha, familiar subordinate, unfamiliar males in dominant and subordinate male mice.** Boxplots show median (horizontal bars), IQR (boxes) and 95% CI (whiskers). Raw data points of each group are also shown on the right side of each box plot.

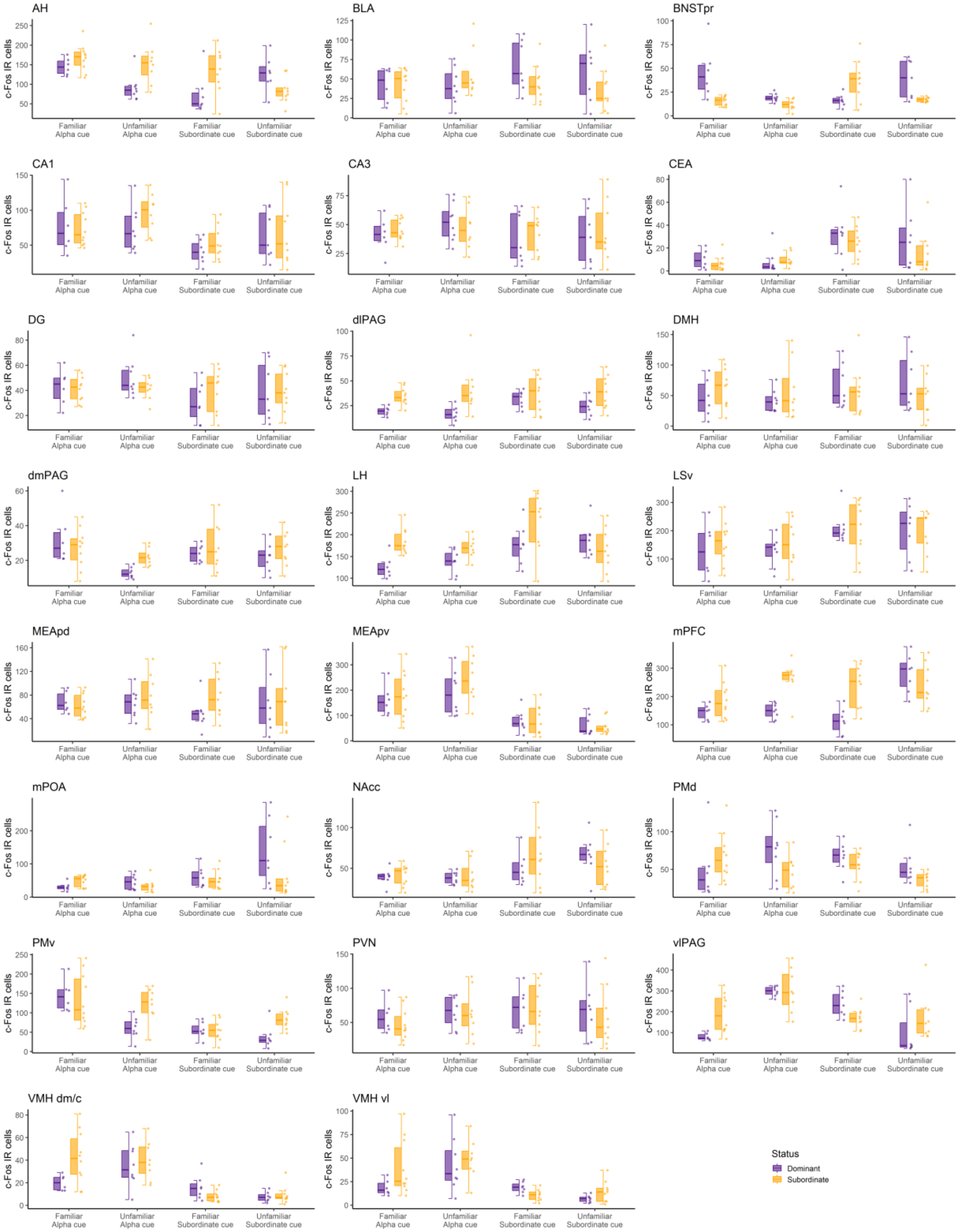

**Supplementary Fig. S3. Result of FDA with cfos count data from SBN nodes (AH, BNSTpr, dlPAG, dmPAG, vlPAG, mPOA, LSv, MEApd, MEApd, VMH vl, VMH dm/c) only.** Function 1 and Function 2 was not as effective as FDA with count data of all brain regions included.

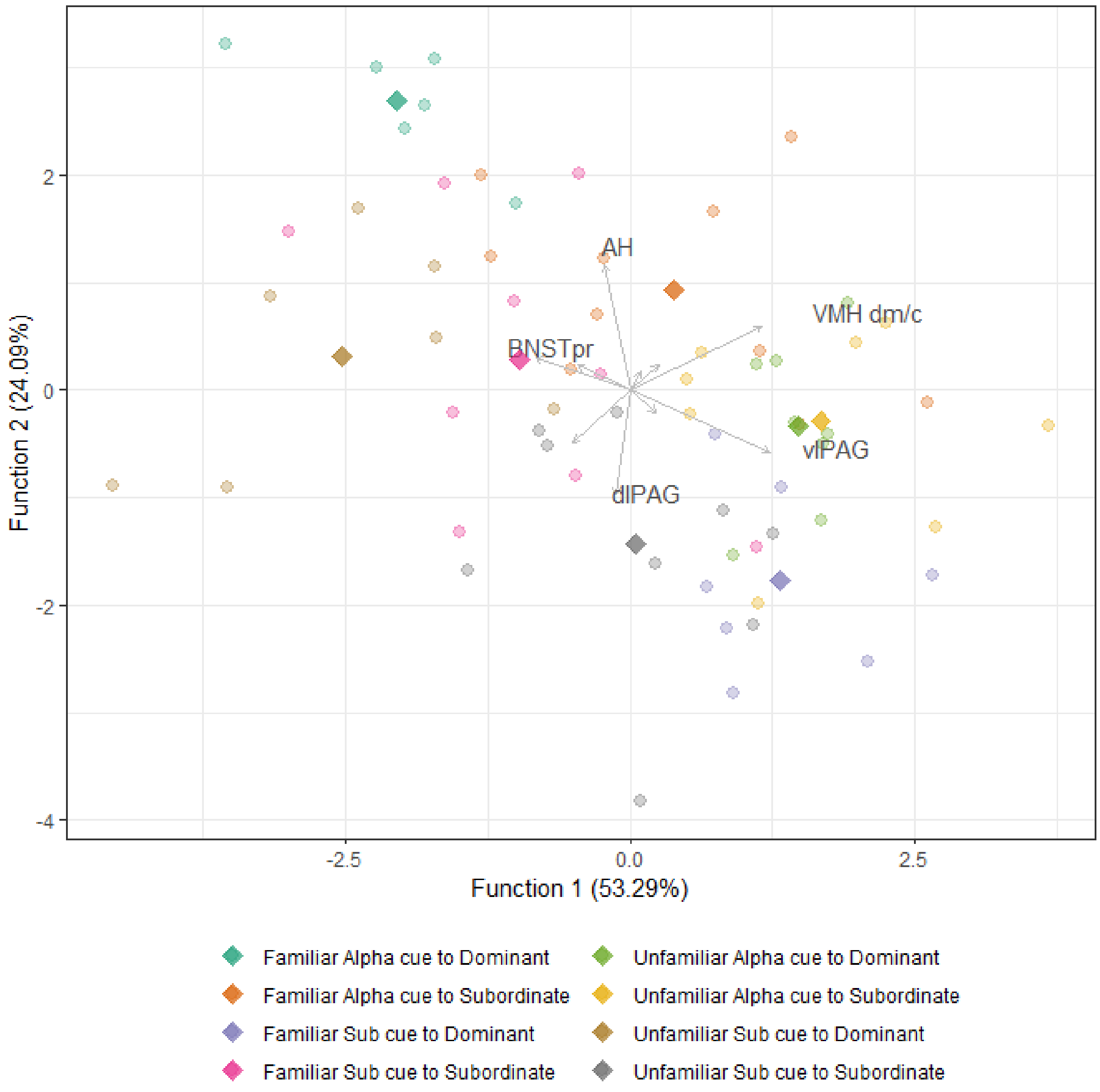

**Supplementary Fig. S4.  Housing Vivarium.**

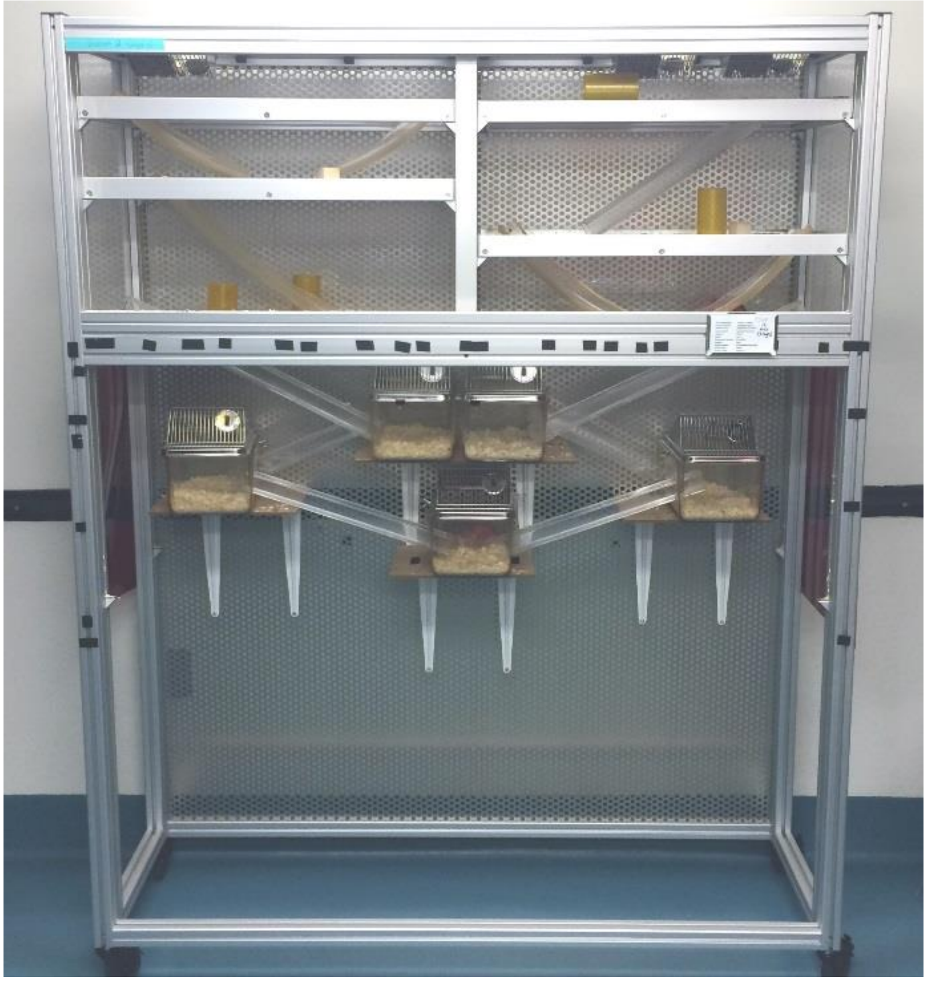

**Supplementary Fig. S5. Sample images of cFOS staining in mPFC, mPOA, DM, VMH dm/c, VMH vl, MEApd, MEApv, PMd, PMv, and vlPAG.**

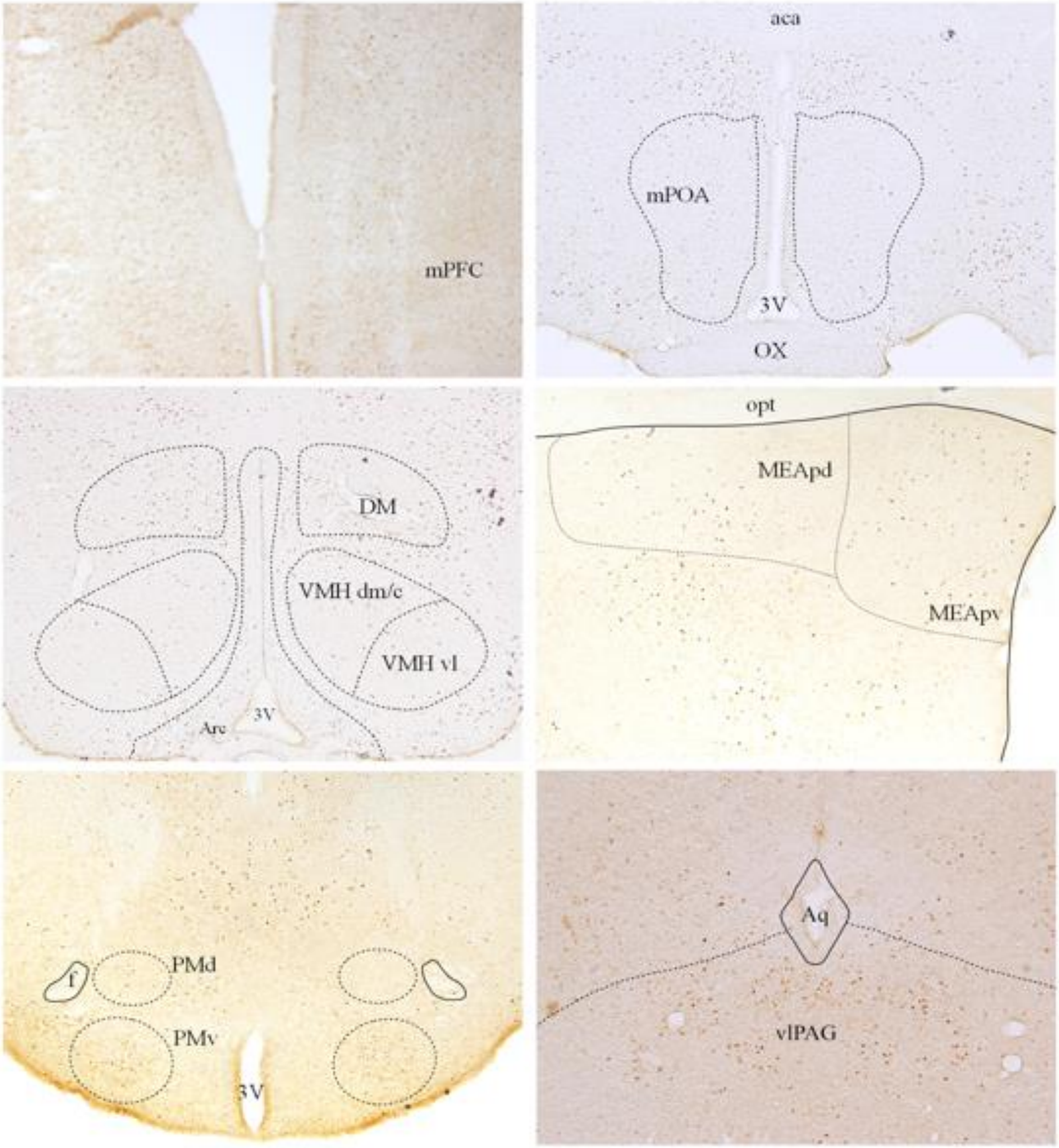

**Supplementary Fig. S6. Individual David's score for each social group**

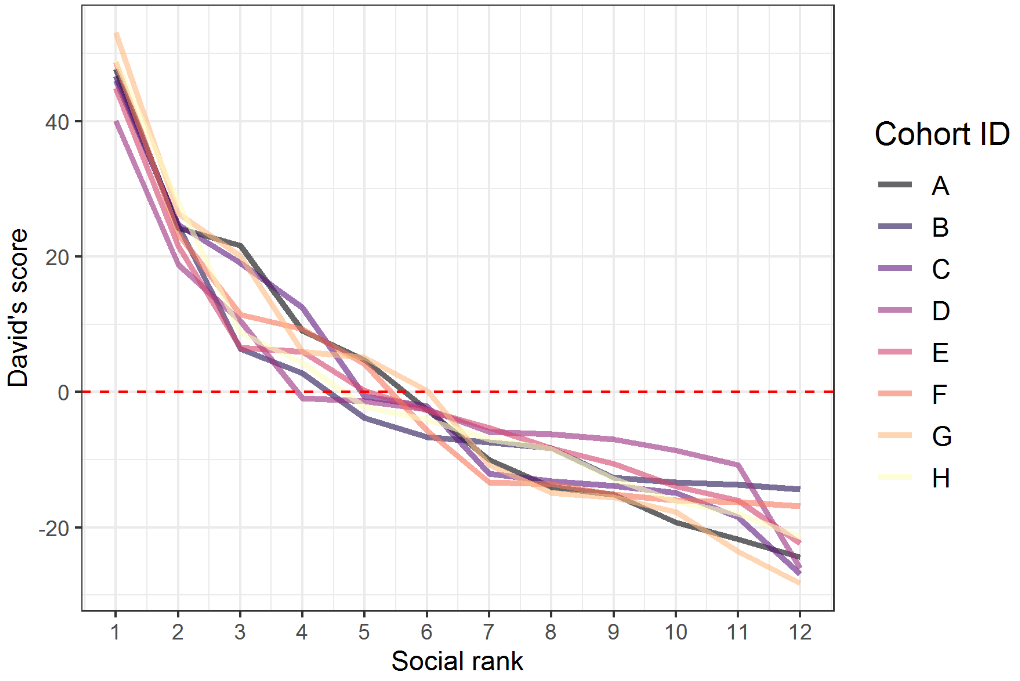
